# Agriculturally sourced multidrug-resistant *Escherichia coli* for use as control strains

**DOI:** 10.1101/2025.03.07.641915

**Authors:** James E. Wells, Lisa M. Durso, Abasiofiok M. Ibekwe, Jonathan G. Frye, Manan Sharma, Clinton F. Williams, Md Shamimuzzaman

**Affiliations:** USDA Agricultural Research Service (ARS), U.S. Meat Animal Research Center, Meat Safety and Quality, Clay Center NE; USDA Agricultural Research Service (ARS), Agroecoystem Management Research, Lincoln, NE; USDA Agricultural Research Service (ARS), Agricultural Water Efficiency and Salinity Research Unit, Riverside CA; USDA Agricultural Research Service (ARS), U.S. National Poultry Research Center, Poultry Microbiological Safety and Processing Research Unit, Athens, GA; USDA Agricultural Research Service (ARS) Environmental Microbial and Food Safety Laboratory, Beltsville, MD; USDA ARS U.S. Arid Land Agricultural Research Center, Water Management and Conservation Research Unit, Maricopa, AZ

**Author notes:** **Corresponding author:** Lisa M. Durso. Mailing address: USDA, ARS, AMRU. Room 251 Keim Hall, UNL-East Campus, Lincoln, NE 68583. **Disclaimer:** Mention of trade names or commercial products in this article is solely for the purpose of providing specific information and does not imply recommendation or endorsement by the U.S. Department of Agriculture. USDA is an equal opportunity provider and employer. **Author contributions: CRediT Conceptualization:** JW, LD, AI, JF, MS, CW **Data curation and formal analysis:** JW, JD, MDS **Methodology:** JW, LD, AI, JF, MS, CW, MDS **Resources:** JW **Validation** JW, LD, AI, JF, MS **Writing-original draft:** LD **Writing – review and edit:** JW, LD, AI, JF, MS, CW, MDS.

**Keywords:** Antibiotic resistance, *Escherichia coli*, Agriculture, Extended-spectrum β-lactamase (ESBL), Tetracycline, Antimicrobial resistance, Control Strains, Surveillance

## Abstract

The environment plays a crucial role in the spread of antibiotic resistance, however environmental surveillance efforts lag far behind clinical surveillance. Natural and human-impacted soil and water are considered major sources of antibiotic resistance in medical and veterinary diagnostics, and the lack of environmental and agricultural surveillance data hinders efforts to assess management impacts or estimate risk. Here we introduce two agriculturally sourced, fully characterized, and genetically sequenced control strains for use in surveillance of extended-spectrum β-lactamase producing (ESBL) and tetracycline-resistant Escherichia coli in the environment, available via publicly accessible culture collections.

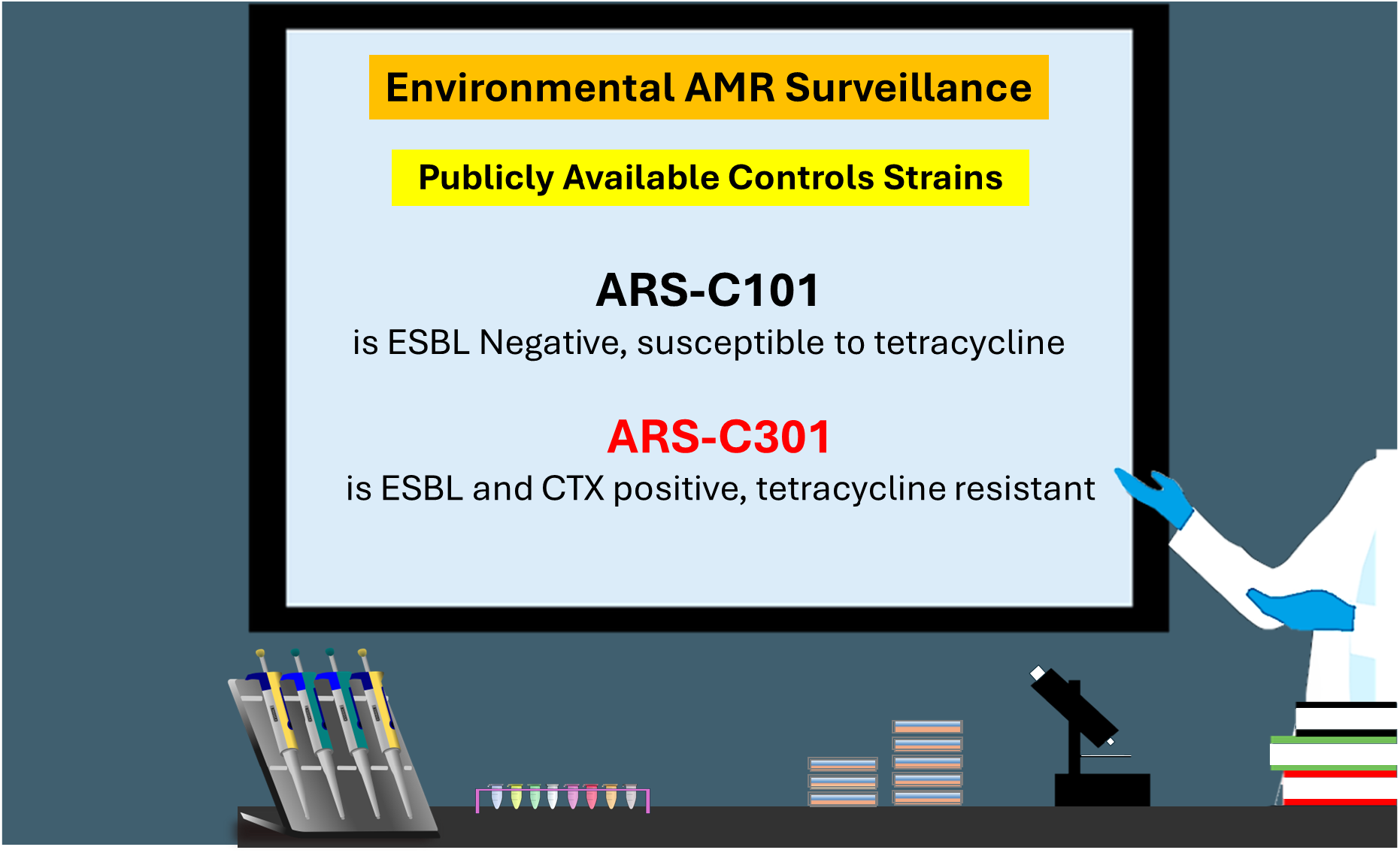

## Body of Text

Antibiotic resistant bacterial infections continue to threaten human and animal health, and critical knowledge gaps remain regarding the role of the environment as a reservoir and transport vehicle for antibiotic resistant bacteria, as well as the potential risks to human, animal, and ecosystem health associated with agricultural and environmental antibiotic resistance (Larsson and Flach, 2022). The global community has embraced a One Health approach for addressing the problem. Recent work highlights the need for standardized methods and quality control measures for environmental monitoring of antibiotic resistance (Liguori et al, 2022). The majority of antibiotic resistant isolates available for use as control strains for methods development, research, and surveillance come from hospital and human public-health laboratories, like the CDC Isolate bank (https://www.cdc.gov/drugresistance/resistance-bank/index.html,2023).

In the context of surveillance and risk assessment, there are important differences between antibiotic resistance in the clinic, and antibiotic resistance in agricultural ecosystems. In the clinic there is a direct, evidence-based connection between the isolate, virulence, and antibiotic resistance. Conversely, most soil-, water-, and airborne bacteria are non-pathogenic. Those that are potential pathogens need to contact and infect a host (person or animal), cause illness, and be non-responsive to the treatment drug before being confirmed as having both the genotypic and phenotypic equivalent of the clinical isolates (Stanton et al., 2022; Hart et al., 2023).

Environmental bacteria have, however, been shown to be the source of clinically important genes encoding resistance to β-lactams, aminoglycosides, amphenicols, sulfonamides, tetracyclines, and other antimicrobials (Forsberg et al., 2012; Hua et al., 2020); and there is a growing awareness that sustainable progress in measuring antibiotic resistance in the clinic is linked to measuring and mitigating it in the natural and agricultural environment.

*Escherichia coli* is increasingly used as a target for environmental surveillance of antibiotic resistance (Anjum et al., 2021; Franklin et al., 2024), and it is well known that human and non-human associated *E. coli* isolates each have different phylogenies, and potentially different virulence profiles (Ochman and Selander, 1984; Denamur et al., 2021). Virulent *E. coli* predominantly fall into groups B2 and D while commensal (non-pathogenic) *E. coli* are generally classified in groups A and B1 (Picard et al., 1999). Even within pathogenic serotypes such as *E. coli* O157:H7, there is a phylogenetic dichotomy between human and non-human isolates, suggesting biological differences in the ability to be transmitted to and/or infect people (Kim et al., 1999; Franz et al, 2019, Ketkhao et al., 2024).

Given these differences, there is a need for broadly accessible non-clinical, non-human associated *E. coli* to serve as controls strains in studies focusing on agricultural and environmental surveillance of antibiotic resistant bacteria and antibiotic resistance method development. To this end, we screened a set of agriculturally associated *E. coli* isolates for their utility as extended spectrum β-lactamase producing (ESBL)/ cefotaxime resistant/tetracycline resistant control strains, in support of One Health efforts. ESBL *E. coli* are a priority target for human health (Hart et al., 2023) and the new National Antibiotic Resistance Monitoring System (NARMS) environmental surveillance effort (Franklin et al., 2024). Tetracycline resistant *E. coli* were chosen as a second target because the tetracyclines are the most widely used antibiotic in food animal production both globally and in the U.S.; and tetracycline is the most commonly studied resistance type in environmental and agricultural samples (Durso and Schmidt, 2017; Mulchandani et al., 2023). The objective was to identify two control strains for use in method development and surveillance: 1) an isolate that displayed phenotypic resistance to tetracycline and cefotaxime, and was positive for ESBL production when evaluated using the Clinical Laboratory Standards Institute combination disk diffusion test (CLSI, 2015); 2) a negative control that was sensitive to both target antibiotics.

Enteric isolates were previously collected from cattle using antibiotic-enriched media, and antibiotic sensitivity testing (AST) was performed for 14 antibiotics from 13 drug classes (Table S1) using the Sensititre™ antimicrobial susceptibility system (TREK Diagnostic Systems Inc., Cleveland, OH, USA) (Long et al., 2023). Of a total of 1133 isolates collected with AST data available, 192 were scored as resistant to cefoxitin, ceftriaxone, or tetracycline, of which 44 were scored as susceptible to all three drugs and 17 were scored as susceptible to all 14 drugs tested.

A subset of twenty-two isolates were chosen for further evaluation and 16 were phenotypically identified as *E. coli* using the indole test, CHROMagar ECC media (DRG International, Springfield, New Jersey, USA), and then confirmed by utilizing the *uidA* PCR specific for *E. coli* (Bej et al., 1991). Confirmed isolates were evaluated for growth on standard and selective media (Tryptic soy Agar (TSA), Tryptone Bile X-glucuronide (TBX) agar), and the selected candidates were tested for phenotype stability through 20 passages on TSA media with and without cefotaxime (4μg/mL) and tetracycline (32 μg/mL).

Isolate ARS C301 was selected as the positive control, and ARS C101 was selected as the negative control (Table 1). Both were genotypically confirmed as *E. coli* via 16S sequencing (Biolog-MIDI Lab, Newark, DE, USA), phylotyped by PCR (Clermont et al., 2000), and subsequently sent to SNPsaurus (Eugene, OR) for whole genome sequencing (Supplementary methods). ARS-C101 and ARS-C301 had 98x and 106x assembled coverage, respectively from 108,488 (ARS-C101) and 130,182 (ARS-C301) reads. Genomes were screened for antibiotic resistance genes using the Comprehensive Antibiotic Resistance Database RGI-Identifier tool (Figure 1). The isolates shared 55 resistance determinants, and ARS-C301 carried an additional 15, including CTX-M-55, LAP-2, and TEM-1 supporting the ESBL phenotype, and *tet*(A) efflux pump supporting the tetracycline resistance phenotype. Additionally, resistance determinants for macrolide, lincosamide, aminoglycoside, fluroquinolone, sulfonamide, rifamycin diaminopyrimidine, and phenicol resistance were also found in the ARS-C301 genome (Supplementary Data File 1). Both strains are publicly available, and the ARS-C301 strain is available as a pre-quantified pellet through Microbiologics (Saint Cloud, MN) (see data availability statement below).

**Table 1.**
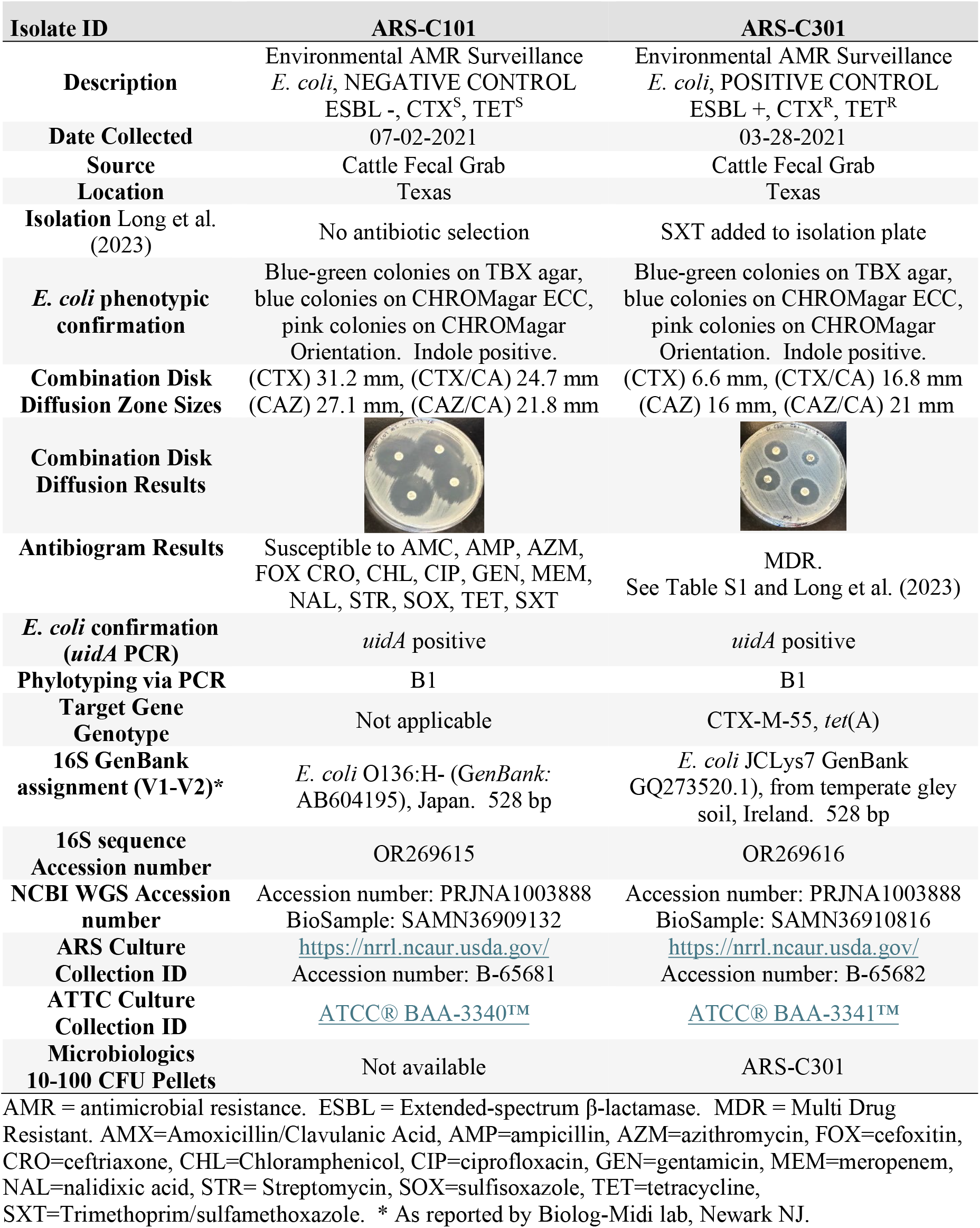
Control Strain Results and Metadata.

| Isolate ID | ARS-C101 | ARS-C301 |
| --- | --- | --- |
| **Description** | Environmental AMR Surveillance<br>*E. coli*, NEGATIVE CONTROL<br>ESBL -, CTX<sup>S</sup>, TET<sup>S</sup> | Environmental AMR Surveillance<br>*E. coli*, POSITIVE CONTROL<br>ESBL +, CTX<sup>R</sup>, TET<sup>R</sup> |
| **Date Collected** | 07-02-2021 | 03-28-2021 |
| **Source** | Cattle Fecal Grab | Cattle Fecal Grab |
| **Location** | Texas | Texas |
| **Isolation** Long et al. (2023) | No antibiotic selection | SXT added to isolation plate |
| ***E. coli* phenotypic confirmation** | Blue-green colonies on TBX agar, blue colonies on CHROMagar ECC, pink colonies on CHROMagar Orientation. Indole positive. | Blue-green colonies on TBX agar, blue colonies on CHROMagar ECC, pink colonies on CHROMagar Orientation. Indole positive. |
| **Combination Disk Diffusion Zone Sizes** | (CTX) 31.2 mm, (CTX/CA) 24.7 mm<br>(CAZ) 27.1 mm, (CAZ/CA) 21.8 mm | (CTX) 6.6 mm, (CTX/CA) 16.8 mm<br>(CAZ) 16 mm, (CAZ/CA) 21 mm |
| **Combination Disk Diffusion Results** |  |  |
| **Antibiogram Results** | Susceptible to AMC, AMP, AZM, FOX CRO, CHL, CIP, GEN, MEM, NAL, STR, SOX, TET, SXT | MDR.<br>See Table S1 and Long et al. (2023) |
| ***E. coli* confirmation (*uidA* PCR)** | *uidA* positive | *uidA* positive |
| **Phylotyping via PCR** | B1 | B1 |
| **Target Gene Genotype** | Not applicable | CTX-M-55, *tet*(A) |
| **16S GenBank assignment (V1-V2)\*** | *E. coli* O136:H- (GenBank: AB604195), Japan. 528 bp | *E. coli* JCLys7 GenBank GQ273520.1), from temperate gley soil, Ireland. 528 bp |
| **16S sequence Accession number** | OR269615 | OR269616 |
| **NCBI WGS Accession number** | Accession number: PRJNA1003888<br>BioSample: SAMN36909132 | Accession number: PRJNA1003888<br>BioSample: SAMN36910816 |
| **ARS Culture Collection ID** | https://nrrl.ncaur.usda.gov/<br>Accession number: B-65681 | https://nrrl.ncaur.usda.gov/<br>Accession number: B-65682 |
| **ATTC Culture Collection ID** | ATCC® BAA-3340™ | ATCC® BAA-3341™ |
| **Microbiologics 10-100 CFU Pellets** | Not available | ARS-C301 |

AMR = antimicrobial resistance. ESBL = Extended-spectrum β-lactamase. MDR = Multi Drug Resistant. AMX=Amoxicillin/Clavulanic Acid, AMP=ampicillin, AZM=azithromycin, FOX=cefoxitin, CRO=ceftriaxone, CHL=Chloramphenicol, CIP=ciprofloxacin, GEN=gentamicin, MEM=meropenem, NAL=nalidixic acid, STR= Streptomycin, SOX=sulfisoxazole, TET=tetracycline, SXT=Trimethoprim/sulfamethoxazole. \* As reported by Biolog-Midi lab, Newark NJ.

**Figure 1.**
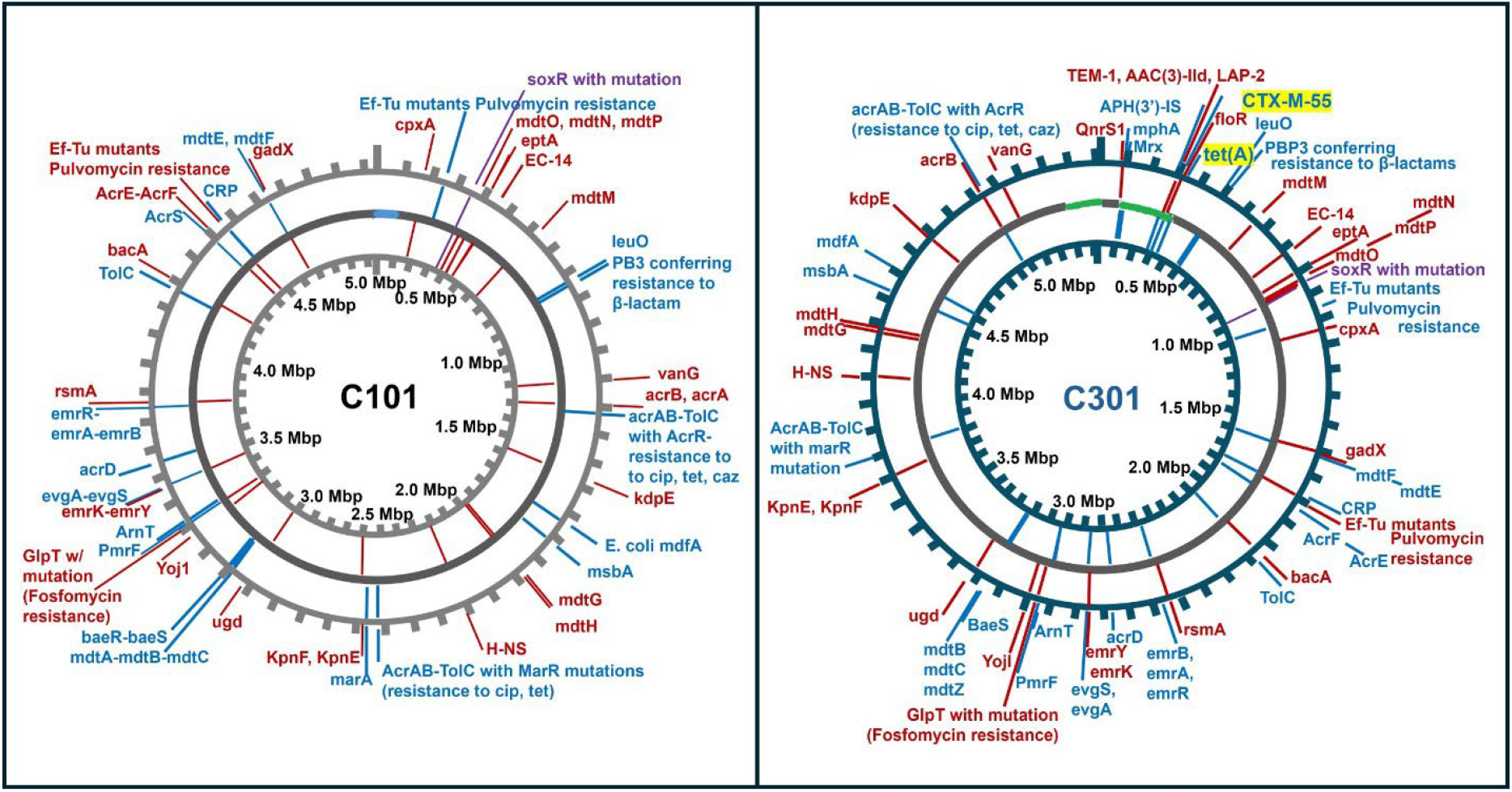
Environmental Control Strain Genome Maps Showing ARGs. ARS-C101 is the negative control strain and has an ESBL negative and tetracycline susceptible phenotype. ARS-C301 is the positive control strain and has an ESBL positive and tetracycline resistant phenotype. The inner- and outer-most circles are for orientation and to indicate gene location. The middle circle represents the genome, with grey indicating regions with full coverage. Genes located on the positive strand are marked in blue, those on the negative strand are marked in red. See also Supplementary Data File 1.

Currently, there is no standardized surveillance of agricultural and environmental antibiotic resistance in the U.S. It is hoped that as new One Health focused efforts are undertaken, the ARS C301 and ARS C101 control strains will be useful in surveillance efforts and method standardization for researchers across the country.

## Data & Bacterial Strain Availability

### ARS-C101 (negative control)

WGS is available from NCBI, accession number PRJNA1003888, BioSample B-65681; Isolate is available from the ARS Culture Collection https://nrrl.ncaur.usda.gov/ (Accession number: B-65681); and from ATCC (isolate ID BAA- 3340).

### ARS-C301 (positive control)

WGS is available from NCBI, accession number PRJNA1003888, BioSample SAMN36910816; Isolate is available from the ARS Culture Collection https://nrrl.ncaur.usda.gov/ (Accession number: B-65682); from ATCC (isolate ID BAA-3341); and from Microbiologics as a quantitative pellet, via a custom order (contact industrial microbiology QC products regional sales manager by email requesting a quote for ARS-C301 https://www.microbiologics.com/contact-us).

## Acknowledgments

We extend thanks to Shannon Ostdiek, Dee Kucera, and Justine Condon for exceptional technical assistance, and to Jasmin Gutierrez, Kassidy Renoe and Antonio Soto for general laboratory support. This work was supported by ARS National Program 212: Soil and Air, and National Program 108 Food Safety.

## SUPPLEMENTARY MATERIALS

### Guide to Supplementary Materials

1. Supplementary Information (This document)
  ▪ Supplementary Methods
  ▪ Supplementary Table 1
  ▪ Reference for Supplementary Information
2. Supplementary Data (Excel document)

## SUPPLEMENTARY METHODS

### Phylotyping

DNA was extracted from isolates using the Qiagen Power Soil kit (Qiagen, Redwood City, CA). Phylotyping was performed as described previously (Ducey et al, 2020), using the Doumith et al. (2012) modification of the Clermont method (Clermont et al., 2000), and the Escobar-Paramo phylotyping scheme (Escobar-Paramo et al., 2004).

### Whole Genome Sequencing

Isolates were grown overnight in tryptic soy broth (Difco, Franklin Lakes, NJ), pelleted and washed twice with phosphate buffered saline, then suspended in DNA Shield (Zymo, Irving, CA) and shipped overnight to Plasmidsaurus (Eugene, OR) for whole genome sequencing using the Nanopore platform. A long-read sequencing library was constructed using Nanopore v14 library prep chemistry (Oxford Nanopore Technologies, Oxford, UK) and run on an Oxford Nanopore R10.4.1 flow cell. The output was assembled using [Flye](https://github.com/fenderglass/Flye) (Kolmogorov et al., 2019) v2.9.1, polished using Nanopore [Medaka](https://github.com/nanoporetech/medaka) v1.8.0, and annotated using [Bakta](https://github.com/oschwengers/bakta) v1.6.1 (Schwengers et al., 2021). Genome wide antibiotic resistance genes (ARGs) were identified using the Comprehensive Antibiotic Resistance Database (CARD) Resistance Gene Identifier (RGI 6.0.3) (Alcock et al., 2023) with default settings and maps were created using Proksee (Grant et al., 2023).

**Supplementary Table 1:** List of antibiotics used in initial screening of isolates.

|  | Drug | Type | ARS-C101<br>(Neg) | ARS-C301<br>(Pos) |
| --- | --- | --- | --- | --- |
| 1 | Amoxicillin/Clavulanic Acid | b-Lactam/b-Lactamase inhibitor<br>combo | S | S |
| 2 | Ampicillin | Penicillins b-lactam | S | R |
| 3 | Azithromycin | Macrolides | S | R |
| 4 | Cefoxitin | Cephens b-lactam | S | S |
| 5 | Ceftriaxone | Cephens b-lactam | S | R |
| 6 | Chloramphenicol | Phenicol | S | R |
| 7 | Ciprofloxacin | Quinolones fluoroquinolone | S | S |
| 8 | Gentamicin | Aminoglycosides | S | R |
| 9 | Meropenem | Penems b-lactam | S | S |
| 10 | Nalidixic Acid | Quinolones | S | S |
| 11 | Streptomycin | Aminoglycosides | S | R |
| 12 | Sulfisoxazole | Folate path antagonists | S | R |
| 13 | Tetracycline | Tetracyclines | S | R |
| 14 | Trimethoprim/Sulfamethoxazole | Folate path antagonists | S | R |

S = susceptible as scored by Sensitre, R = resistant as scored by Sensitre. Results collected and reported by Long et al. (2023) https://doi.org/10.1089/fpd.2023.0001, using methods from Agga et al. (2016) https://doi.org/10.1089/mdr.2015.0259

## Notes

https://doi.org/10.15482/USDA.ADC/27138789.v1

### Competing Interest Statement

The authors have declared no competing interest.

## References

1. Anjum, M.F., Schmitt, H., Börjesson, S., Berendonk, T.U., Donner, E., Stehling, E.G., Boerlin, P., Topp, E., Jardine, C., Li, X. and Li, B., 2021. The potential of using E. coli as an indicator for the surveillance of antimicrobial resistance (AMR) in the environment. Current Opinion in Microbiology, 64, pp.152–158.

2. Bej AK, DiCesare JL, Haff JL, Atlas RM (1991) Detection of Escherichia coli and Shigella spp. in water by using the polymerase chain reaction and gene probes for uid. Appl Environ Microbiol 57: 1013–1017.

3. Clermont, O., Bonacorsi, S. and Bingen, E., 2000. Rapid and simple determination of the Escherichia coli phylogenetic group. Applied and environmental microbiology, 66(10), pp.4555–4558.

4. CLSI. (2015). Performance Standards for Antimicrobial Susceptibility Testing; Twenty-Fifth Informational Supplement. Wayne, PA: Clinical and Laboratory Standards Institute.

5. Denamur, E., Clermont, O., Bonacorsi, S. and Gordon, D., 2021. The population genetics of pathogenic Escherichia coli. Nature Reviews Microbiology, 19(1), pp.37–54.

6. Durso, L.M. and Schmidt, A.M., 2017. Antimicrobial resistance related to agricultural wastewater and biosolids. Antimicrobial Resistance in Wastewater Treatment Processes, pp.219–240.

7. CDC & FDA Antibiotic Resistance Isolate Bank | CDC https://www.cdc.gov/drugresistance/resistance-bank/index.html

8. Forsberg KJ, Reyes A, Wang B, Selleck EM, Sommer MO, Dantas G. The shared antibiotic resistome of soil bacteria and human pathogens. Science. 2012 Aug 31;337(6098):1107–11. doi: 10.1126/science.1220761. PMID: 22936781; PMCID: PMC4070369.

9. Franklin, A.M., Weller, D.L., Durso, L.M., Bagley, M., Davis, B.C., Frye, J.G., Grim, C.J., Ibekwe, A.M., Jahne, M.A., Keely, S.P. and Kraft, A.L., 2024. A one health approach for monitoring antimicrobial resistance: developing a national freshwater pilot effort. Frontiers in water, 6, p.1359109.

10. Franz, E., Rotariu, O., Lopes, B.S., MacRae, M., Bono, J.L., Laing, C., Gannon, V., Söderlund, R., Van Hoek, A.H., Friesema, I. and French, N.P., 2019. Phylogeographic analysis reveals multiple international transmission events have driven the global emergence of Escherichia coli O157: H7. Clinical Infectious Diseases, 69(3), pp.428–437.

11. Hart, A., Warren, J., Wilkinson, H. and Schmidt, W., 2023. Environmental surveillance of antimicrobial resistance (AMR), perspectives from a national environmental regulator in 2023. Eurosurveillance, 28(11), p.2200367.

12. Hua, M., Huang, W., Chen, A., Rehmet, M., Jin, C. and Huang, Z., 2020. Comparison of antimicrobial resistance detected in environmental and clinical isolates from historical data for the US. BioMed research international, 2020.

13. Jacob, Megan E., Shivaramu Keelara, Awa Aidara-Kane, Jorge R. Matheu Alvarez, and Paula J. Fedorka-Cray. “Optimizing a screening protocol for potential extended-spectrum β-lactamase Escherichia coli on MacConkey agar for use in a global surveillance program.” Journal of Clinical Microbiology 58, no. 9 (2020): e01039–19.

14. Karp, B.E., Tate, H., Plumblee, J.R., Dessai, U., Whichard, J.M., Thacker, E.L., Hale, K.R., Wilson, W., Friedman, C.R., Griffin, P.M. and McDermott, P.F., 2017. National antimicrobial resistance monitoring system: two decades of advancing public health through integrated surveillance of antimicrobial resistance. Foodborne pathogens and disease, 14(10), pp.545–557.

15. Ketkhao, P., Utrarachkij, F., Parikumsil, N., Poonchareon, K., Kerdsin, A., Ekchariyawat, P., Narongpun, P., Nakajima, C., Suzuki, Y. and Suthienkul, O., 2024. Phylogenetic diversity and virulence gene characteristics of Escherichia coli from pork and patients with urinary tract infections in Thailand. Plos one, 19(7), p.e0307544.

16. Kim, J., Nietfeldt, J. and Benson, A.K., 1999. Octamer-based genome scanning distinguishes a unique subpopulation of Escherichia coli O157: H7 strains in cattle. Proceedings of the National Academy of Sciences, 96(23), pp.13288–13293.

17. Kolmogorov, M., Yuan, J., Lin, Y., and Pevzner, P., 2019. Assembly of long error-prone reads using repeat graphs. Nature Biotechnology, doi:10.1038/s41587-019-0072-8.

18. Larsson, D.J. and Flach, C.F., 2022. Antibiotic resistance in the environment. Nature Reviews Microbiology, 20(5), pp.257–269.

19. Liguori, K., Keenum, I., Davis, B.C., Calarco, J., Milligan, E., Harwood, V.J. and Pruden, A., 2022. Antimicrobial resistance monitoring of water environments: a framework for standardized methods and quality control. Environmental science & technology, 56(13), pp.9149–9160.

20. Long, N.S., Hales, K.E., Berry, E.D., Legako, J.F., Woerner, D.R., Broadway, P.R., Carroll, J.A., Burdick Sanchez, N.C., Fernando, S.C. and Wells, J.E., 2023. Antimicrobial Susceptibility of Trimethoprim–Sulfamethoxazole and 3rd-Generation Cephalosporin-Resistant Escherichia coli Isolates Enumerated Longitudinally from Feedlot Arrival to Harvest in High-Risk Beef Cattle Administered Common Metaphylactic Antimicrobials. Foodborne Pathogens and Disease.

21. Mulchandani, R., Wang, Y., Gilbert, M. and Van Boeckel, T.P., 2023. Global trends in antimicrobial use in food-producing animals: 2020 to 2030. PLOS Global Public Health, 3(2), p.e0001305.

22. Ochman, H. and Selander, R.K., 1984. Standard reference strains of Escherichia coli from natural populations. Journal of bacteriology, 157(2), pp.690–693.

23. Picard, B., Garcia, J.S., Gouriou, S., Duriez, P., Brahimi, N., Bingen, E., Elion, J. and Denamur, E., 1999. The link between phylogeny and virulence in Escherichia coli extraintestinal infection. Infection and immunity, 67(2), pp.546–553.

24. Stanton, I.C., Bethel, A., Leonard, A.F.C., Gaze, W.H. and Garside, R., 2022. Existing evidence on antibiotic resistance exposure and transmission to humans from the environment: a systematic map. Environmental evidence, 11(1), pp.1–24.

25. Schwengers O., Jelonek L., Dieckmann M. A., Beyvers S., Blom J., Goesmann A., 2021. Bakta: rapid and standardized annotation of bacterial genomes via alignment-free sequence identification. Microbial Genomics, 7(11). 10.1099/mgen.0.000685.

26. World Health Organization, 2021. “WHO integrated global surveillance on ESBL-producing E. coli using a “One Health” approach: implementation and opportunities.”. ISBN: 978-92-4-002140-2 https://www.who.int/publications-detail-redirect/9789240021402

## SUPPLEMENTARY INFORMATION REFERENCES

Agga, G.E., Schmidt, J.W. and Arthur, T.M., 2016. Antimicrobial-resistant fecal bacteria from ceftiofur-treated and nonantimicrobial-treated comingled beef cows at a cow–calf operation. Microbial drug resistance, 22(7), pp.598–608. 10.1089/mdr.2015.0259

Alcock, B.P.; Huynh, W.; Chalil, R.; Smith, K.W.; Raphenya, A.R.; Wlodarski, M.A.; Edalatmand, A.; Petkau, A.; Syed, S.A.; Tsang, K.K.; et al. CARD 2023: Expanded Curation, Support for Machine Learning, and Resistome Prediction at the Comprehensive Antibiotic Resistance Database. Nucleic Acids Res. 2023, 51, D690–D699.

Clermont, O., Bonacorsi, S. and Bingen, E., 2000. Rapid and simple determination of the Escherichia coli phylogenetic group. Applied and environmental microbiology, 66(10), pp.4555–4558.

Doumith, M., Day, M.J., Hope, R., Wain, J. and Woodford, N., 2012. Improved multiplex PCR strategy for rapid assignment of the four major Escherichia coli phylogenetic groups. Journal of clinical microbiology, 50(9), pp.3108–3110.

Ducey, T.F., Durso, L.M., Ibekwe, A.M., Dungan, R.S., Jackson, C.R., Frye, J.G., Castleberry, B.L., Rashash, D.M., Rothrock, M.J., Boykin, D. and Whitehead, T.R., 2020. A newly developed Escherichia coli isolate panel from a cross section of US animal production systems reveals geographic and commodity-based differences in antibiotic resistance gene carriage. Journal of hazardous materials, 382, p.120991.

Escobar-Páramo, P., Grenet, K., Le Menac’h, A., Rode, L., Salgado, E., Amorin, C., Gouriou, S., Picard, B., Rahimy, M.C., Andremont, A. and Denamur, E., 2004. Large-scale population structure of human commensal Escherichia coli isolates. Applied and environmental microbiology, 70(9), pp.5698–5700.

Grant, J. R., Enns, E., Marinier, E., Mandal, A., Herman, E. K., Chen, C. Y., et al. (2023). Proksee: in-depth characterization and visualization of bacterial genomes. Nucleic Acids Res. gkad326. doi: 10.1093/nar/gkad326

Kolmogorov, M., Yuan, J., Lin, Y., and Pevzner, P., 2019. Assembly of long error-prone reads using repeat graphs. Nature Biotechnology, doi:10.1038/s41587-019-0072-8.

Schwengers O., Jelonek L., Dieckmann M. A., Beyvers S., Blom J., Goesmann A., 2021. Bakta: rapid and standardized annotation of bacterial genomes via alignment-free sequence identification. Microbial Genomics, 7(11). https://doi.org/10.

